# Growth and patterning in vertebrate limb development A timescale perspective on skeletal specification

**DOI:** 10.1101/2025.03.20.644440

**Authors:** S Ben Tahar, E Comellas, TJ Duerr, D Bakr, JR Monaghan, JJ Muñoz, SJ Shefelbine

**Affiliations:** Northeastern University; Universitat Politècnica de Catalunya

## Abstract

1

The vertebrate limb provides a powerful system to study how growth and molecular signaling interact to shape complex skeletal patterns. However, how these processes are coordinated across space and time is not fully understood. This study introduces a computational tool to examine how growth interacts with positional cues and self-organizing patterning mechanisms to shape skeletal structures in both mice and axolotl limbs. We developed the Growth-Processing-Propagation (GPP) framework, a reaction-diffusion system within a growing domain, in which the relative contribution of growth, processing (reaction) and propagation (diffusion) is modulated through two non-dimensional parameters, informed by experimental morphogen maps. This formulation normalizes the reaction-diffusion equation relative to growth, enabling investigation of how different spatiotemporal regimes of growth, processing and propagation interact to produce whole limb patterning. The GPP framework captures the progressive formation of limb segments and digit patterns by varying the relative contributions of reaction, diffusion, and growth to the pattern. Our models indicate that in the proximal region (humerus, radius/ulna) the contributions of growth, reaction and diffusion are equally important to patterning, but in the distal elements (hands) the reaction and diffusion contributions are much greater than the contribution of growth to formation of the digits. A single framework predicts whole-limb skeletal patterns in both mice and axolotls, despite their morphological differences, highlighting its potential to explore conserved and divergent features of limb development from an evolutionary perspective through a unified mechanism across species.

Significance
Understanding limb skeletal patterning is a central question in developmental biology. Numerical models provide a means to explore this process. We introduce the Growth-Processing-Propagation (GPP) framework, which integrates tissue expansion (growth), gene regulation and cellular specification (processing), and signaling spread (propagation). By normalizing patterning dynamics relative to growth, the framework reveals how the relative contributions of growth, processing, and propagation vary across the limb, influencing the timing and positioning of skeletal elements. The model successfully predicts whole-limb skeletal patterning in both mice and axolotls, supporting the idea of a shared underlying mechanism. This cross-species framework offers a versatile computational tool for studying fundamental processes in morphogenesis, with potential applications in developmental biology, and evolutionary studies.

## 2 Introduction

Tetrapod limb development is an exemplar model for exploring how developmental processes are coordinated during organogenesis. The developing limb is patterned along three primary axes: the proximodistal axis (PD), which is segmented into stylopod (humerus or femur), zeugopod (radius/ulna or tibia/fibula), and autopod (hand or foot with digits), the anteroposterior axis (AP), which specifies the number of digits [1], and the dorsoventral axis (DV). Numerical simulations of limb patterning have served as essential exploratory tools for understanding how self-organization and tissue growth dynamics give rise to complex skeletal structures. They are primarily based on two complementary theoretical frameworks: positional information and reaction-diffusion (Turing) systems.

Two well-characterized signaling centers during limb development are the Apical Ectodermal Ridge (AER), which provides distal cues guiding PD outgrowth and specification [2, 3], and the Zone of Polarizing Activity (ZPA), which controls AP identity and growth [4, 1]. Past studies suggest that these two centers do not function independently, but instead are part of a feedback loop to coordinate patterning across both axes [5, 6]. Both the AER or the ZPA have been proposed as sources of morphogens gradients, which are central to the positional information (PI) framework [7, 3, 8, 9], which states that diffusible molecules, known as morphogens, form gradients that establish a coordinate system within the limb bud, giving positional identity to cells [4]. Cells interpret specific morphogen concentrations to acquire their distinct fates. The discovery of *Hox* genes provided a molecular basis for this idea, with *Hoxa9* expressed throughout the limb, *Hoxa11* in the zeugopod and autopod, and *Hoxa13* in the autopod [10]. These genes are activated in response to a morphogen gradient, such as retinoic acid from the flank [11, 12, 13]. The conservation of these expression patterns across tetrapods suggests that positional information acts as a shared blueprint for limb development that is sufficiently flexible to allow for variations in limb morphology [14, 15]. While this model has been instrumental in explaining broad patterning mechanisms, it primarily addresses how cells acquire positional identity, rather than how distinct skeletal elements (such as humerus, radius, ulna or digits) emerge as spatially organized patterns.

In response to these limitations, there has been a renewed focus on Turing systems, or reaction-diffusion (RD) models, as a mechanism to produce self-organized spatial patterns [16]. These models describe the interactions between two or more morphogens through coupled partial differential equations generating stable molecular patterns that guide cell fate. Understanding digit periodicity using Turing-based numerical models has provided foundational insights into limb patterning [17, 18, 19, 20]. Significant advances have been made in identifying the gene regulatory networks driving these dynamics [21, 17], while simultaneously revealing their inherent complexity. Recent studies have highlighted that these networks are highly context-dependent, with the same molecular components performing different functions based on spatial location and developmental timing [5, 22, 23]. Similarly, our understanding of morphogen movement has evolved beyond simple diffusion models, with experimental evidence showing that receptor-mediated interactions significantly influence morphogen transport [24]. These findings suggest an opportunity to conceptualize RD systems in terms of functional interactions rather than specific molecular identities. Building on this functional perspective, tissue growth adds an additional layer of complexity that shapes when, where, and how patterns emerge.

Growth is the continuous change in limb shape and size throughout development which occurs through cell proliferation and matrix production. Geometry (i.e. template shape) influences the patterns generated by reaction-diffusion models [25]. In addition to accurately capturing changes in geometry during growth, the model must account for how growth transports molecular signals within the tissue. Mathematically, this is translated into a convective term in the reaction-diffusion system, which depends on the growth rate [26]. Both aspects, growth of the template and growth-mediated transport of molecules, are needed to model patterning within a growing domain. Previous computational models simulating limb development have incorporated growth through different approaches. Some studies solve equations in idealized, synthetic domains to isolate specific growth effects [27, 28, 29, 30]. Complementary to these, significant contributions have been made in characterizing growth dynamics and tissue movement in realistic biological geometries, particularly in the mouse limb bud [31, 32, 33, 34]. Our work aims to bridge these approaches by implementing the convective term derived from mathematical growth models in accurate limb geometries across developmental stages.

In this paper, we present a framework that integrates experimentally measured shape profiles with the Turing system and positional information. Building on established models [35, 21, 18], we propose that positional information regulates the parameters controlling the relative contribution of reaction and diffusion with respect to growth during pattern formation. This perspective allows us to explore reaction-diffusion systems as processing-propagation events, and to quantify how the strength of these events relative to growth varies across different regions of the growing domain. By modulating the weight of reaction and diffusion terms in our equations with respect to growth, our model captures the continuous emergence of different limb segments in both mice and axolotls.

## 3 Results

### 3.1 The Growth-Processing-Propagation model

We propose a two species reaction-diffusion (RD) equation that models the growth and differentiation of the limb skeletal elements. Within this species framework, the species *u* represents the specification of the skeletal elements (i.e. differentiation to cartilage), while *v* represents an antagonist of this specification. As a result, the patterns generated in a forelimb model correspond to skeletal elements: the humerus, radius, ulna, and digits.

The RD model within a growing domain is interpreted as a Growth-Processing-Propagation (GPP) framework (Fig. 1A). Processing (represented by the reaction term) refers to the cellular mechanisms by which specification information is interpreted and acted upon. This may include transcriptional activation of genes, signal transduction cascades, or other intracellular processes that regulate cell fate decisions. Propagation (represented by the diffusion term) refers to the transmission of specification information across the tissue. This can occur through the diffusion of signaling molecules or through direct cell-cell communication mechanisms, such as gap junctions or mechanical coupling. Growth in this context involves both cell proliferation and extracellular matrix production, which drive the shape changes of the limb bud throughout development. The GPP framework incorporates growth in three ways: through an evolving geometry as model input, through a convective term in the equations, and through non-dimensionalization with respect to the growth timescale. The convective term accounts for signal transport caused by tissue expansion and is dependent on growth rate. The amount of growth is determined from images of limb bud profiles through development [36] (see Materials and Methods). Incorporating growth directly in the equation and numerically in the computational domain provides a framework to account for how growth rate affects pattern formation. Additionally, this integration produces a model where processing and propagation interact with growth through two parameters, *α*_*R*_ and *β*_*D*_, to create patterns.

**Figure 1.**
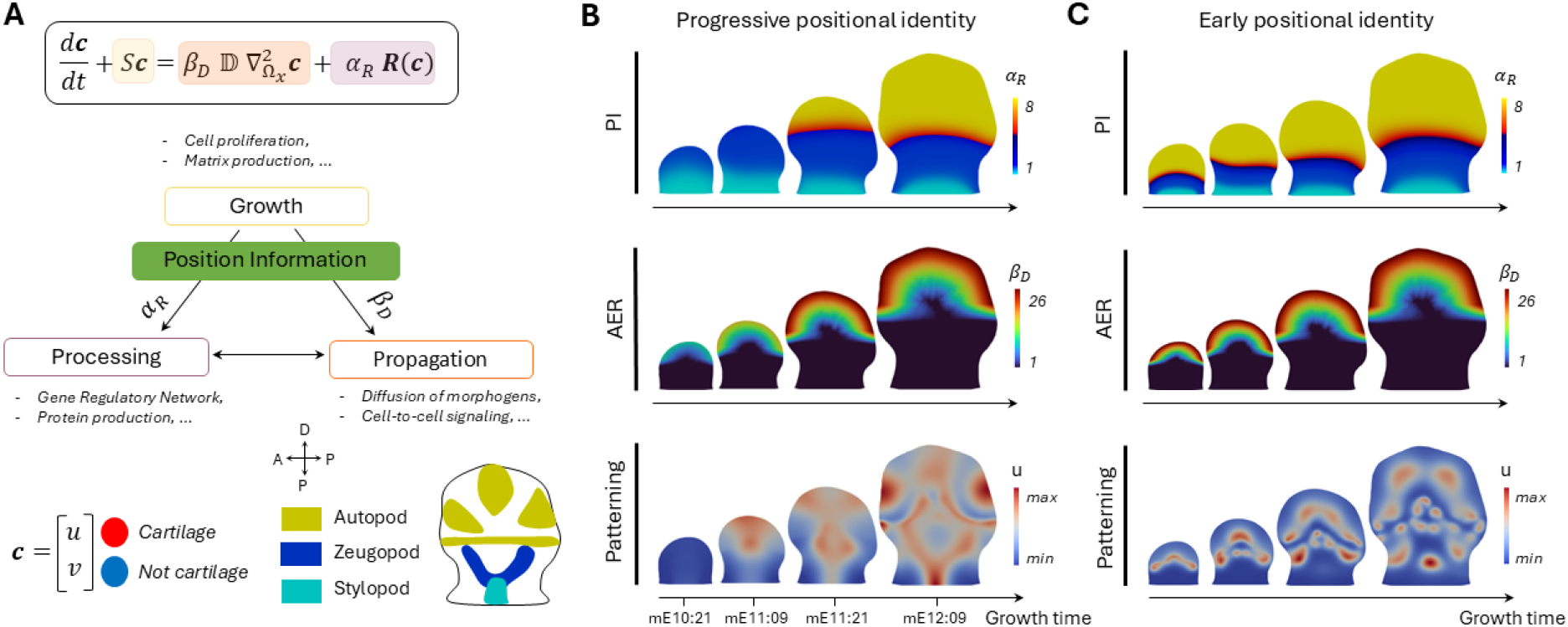
Skeletal patterning of a representative growing mouse limb bud predicted by the GPP computational framework. (**A**) **c** is a vector of the two out-of-phase species, *u* and *v*, where high values of *u* indicate differentiation of skeletal elements through cartilage condensation. The convective term *S***c** captures the effect of growth, with *S* representing the growth rate. The diffusion term 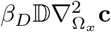 represents propagation, and the reaction term *α*_*R*_**R**(**c**) represents processing. *α*_*R*_ and *β*_*D*_ indicate the contribution of reaction and diffusion relative to growth in the system. Growth, processing and propagation refer to tissue expansion, gene regulation and signaling communication. (**B**) The first row shows the distribution of *α*_*R*_, informed by proximodistal Positional Information (PI) patterns. The second row displays the input map distribution of *β*_*D*_, informed by signaling from the Apical Ectodermal Ridge (AER). The final row shows the predicted skeletal patterning, with regions of high *u* values shown in red indicating locations of cartilage condensation. The humerus, followed by the radius and ulna, and, finally, five digits can be identified. (**C**) The distribution of *α*_*R*_ and *β*_*D*_ are changed to represent early positional identity. This results in incomplete patterning. Our model suggests that the identity of the three limb segments (stylopod, zeugopod, and autopod) occurs sequentially (Fig. 1B) and is not predetermined at the limb bud stage.

Mathematically, we describe the GPP system with a non-dimensionalized version of the RD equation in a growing domain [27],

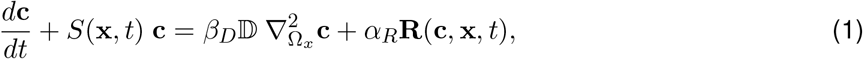

where *d***c***/dt* is the total time derivative of the species concentration **c** = [*u, v*], *S*(**x**, *t*) **c** is the convective term accounting for how growth influences the RD system with *S* being the growth rate, 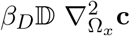 represents propagation through a diffusion process, and *α*_*R*_**R**(**c, x**, *t*) corresponds to processing through the reaction term (see Materials and Methods). The parameters *α*_*R*_ and *β*_*D*_ represent the relative times-cales of growth to processing (reaction) and propagation (diffusion), respectively, and vary across time and space of the growing domain. Values of these parameters below 1 indicate that growth contributes more significantly to pattern formation than processing or propagation, while values above 1 reflect a greater contribution from processing and propagation relative to growth. See Section Materials and Methods for the expression of *α*_*R*_ and *β*_*D*_ as a function of the characteristic times.

The GPP framework builds on previous efforts to combine Turing’s reaction-diffusion theory with Wolpert’s positional information (PI) [35, 21, 18]. PI acts upstream of the GPP system, modulating the two key parameters that control the relative contributions of growth, processing, and propagation in pattern formation, i.e. *α*_*R*_ increasing along the proximodistal axis and *β*_*D*_ exhibiting higher values in distal regions. As the limb bud grows and changes shape over time, these spatial variations in *α*_*R*_ and *β*_*D*_ become more pronounced. The parameter *α*_*R*_ controls the wavelength of the pattern, which determines the number of stripes (skeletal elements) formed. As *α*_*R*_ increases distally, the wavelength shortens, allowing a transition from one stripe (humerus) to two stripes (radius and ulna) and eventually to multiple stripes (digits). An increase in *α*_*R*_ from proximal to distal suggests that processing contributes more to patterning, relative to growth, in distal segments. The parameter *β*_*D*_ governs the orientation of the stripe-like patterns, by controlling the relative weight of propagation to growth at the distal tip of the limb bud. While *α*_*R*_ and *β*_*D*_ represent the relative contributions of processing and propagation compared to growth, their spatial profiles are inspired by developmental trends: *α*_*R*_ reflects the proximodistal PI conveyed by *Hoxa* gene expression [37], and *β*_*D*_ captures aspects of signaling associated with the AER [38]. Though not direct representations of gene expression or tissue structures, these parameters act as spatial modulations within the model that align with known biological patterning influences.

The GPP framework provides a tool to investigate how the temporal evolution of positional identity affect patterning outcomes, allowing us to compare two classic theoretical models for the establishment of positional identity: progressive positional identity (Fig. 1B) and early positional identity (Fig. 1C) [39]. The progressive model suggests that positional identity is acquired sequentially, resulting in the stylopod determined first, followed by the zeugopod, and finally the autopod [40]. In contrast, in the early model (Fig. 1C), distinct regions of positional identity are assigned at the limb bud stage and expand with tissue growth. To simulate these two scenarios, we varied the temporal progression of the spatial input maps *α*_*R*_ and *β*_*D*_. In the progressive identity scenario, *α*_*R*_ starts as a single uniform value and gradually differentiates into two and then three distinct regions, each with a different value, corresponding to the limb segments. Similarly, *β*_*D*_ values increase over time, especially in distal regions. In the early identity scenario, both *α*_*R*_ and *β*_*D*_ distributions and values are fixed from the limb bud stage and are stretched with the domain growth. Only the progressive identity scenario leads to appropriate skeletal patterning, with sequential emergence of the humerus, radius/ulna, and digits. These results support a model in which positional identity unfolds progressively during development, rather than being fully established at early stages.

To explore how positional cues shape skeletal patterning, we systematically modified the presence of *α*_*R*_ and *β*_*D*_ in the model (Fig. 2). In the absence of both PI and AER, implemented as constant and homogeneous distributions of *α*_*R*_ and *β*_*D*_, the framework predicts simple stripe patterns, potentially corresponding to a rudimentary stylopod element (Fig. 2A1). The absence of AER, modeled as a constant and homogeneous *β*_*D*_ while allowing *α*_*R*_ to vary as in Fig. reffig:GPP, disrupts stripe orientation (Fig. 2A2). It highlights the critical role of the AER in influencing digit patterning [41, 42]. Including AER but excluding PI, by using a constant and homogeneous *α*_*R*_ and keeping the variations in *β*_*D*_ as in Fig. reffig:GPP, does not produce a stripe-like pattern for more proximal skeletal elements (Fig. 2A3).

**Figure 2.**
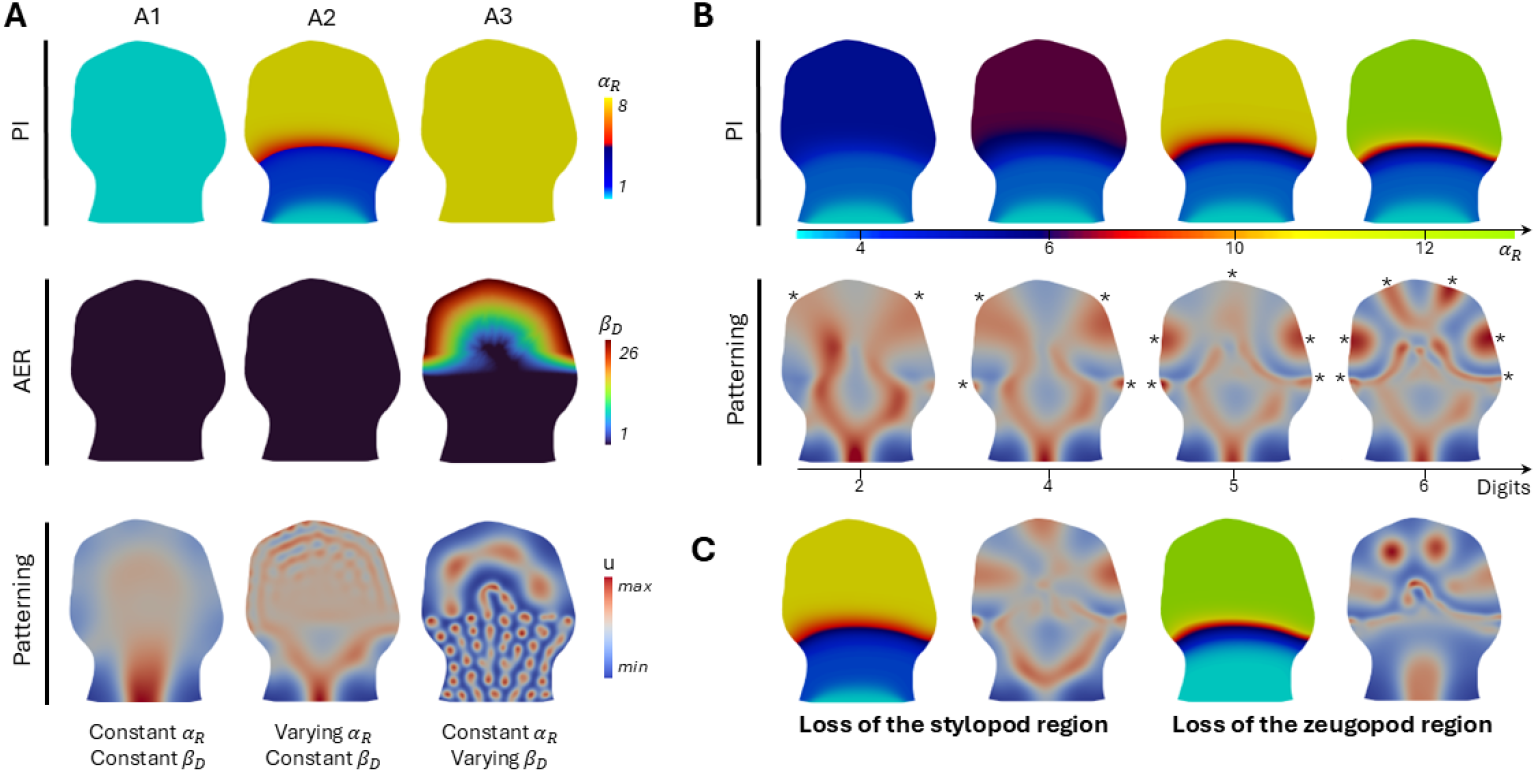
Modifying positional cues and predicting limb segment outcomes with the GPP framework. The final PI, AER and skeletal pattern distribution is represented. (**A**) A1 represents the framework without incorporating PI or AER, implemented as constant and homogeneous *α*_*R*_ and *β*_*D*_, resulting in a simple stripe pattern. A2 includes PI by allowing spatial variation in *α*_*R*_ but excludes AER by keeping *β*_*D*_ constant and homogeneous. This preserves the proximal pattern seen in Fig. 1B but disrupts stripe orientation. A3 incorporates AER by using a spatially varying *β*_*D*_ but excludes PI with constant *α*_*R*_, leading to the loss of more proximal skeletal elements. (**B**) Increasing the strength of processing relative to growth (controlled by the parameter *α*_*R*_) in the autopod region leads to a greater number of digits. The stars indicates the digits. (**C**) Modifying the spatial boundaries of the stylopod or zeugopod regions using the values of *α*_*R*_ can result in the loss of proximal skeletal limb segments.

Furthermore, in the GPP framework, PI is defined along the proximal-distal axis through the spatial modulation of *α*_*R*_, yet the resulting pattern also reflects anterior-posterior organization. By adjusting the weight of processing relative to growth, we can modulate the number of digits formed, ranging from two to six (Fig. 2B). This highlights how PD inputs can influence AP patterning outcomes and reinforces the known interdependence between the two axes [43, 44].

The framework cannot completely remove the stylopod or zeugopod without altering autopod patterning (Fig. 2C). However, by shifting the transition points between different *α*_*R*_ values (cyan, blue and yellow regions in Fig. 2C), we can shorten specific segments, in accordance with biological experiments [45].

### 3.2 Integrating experimental data to predict axolotl limb patterning

To test the robustness of the GPP framework across divergent tetrapod species, we applied it to the axolotl salamander (Ambystoma mexicanum), an anamniote with well-documented differences in limb development compared to amniotes such as mice and chickens [46]. While several features of their development are unique, key principles are conserved [47, 48, 49, 50].

One of the most striking differences is that salamanders do not form a morphologically distinct AER. Without forming a thickened pseudostratified epithelium, the region retain its function to give distal cues [49]. Molecular signaling pathways also diverge in some aspects. For instance, while FGF8 is restricted to the AER in mice and chickens, in axolotls it is only expressed in the mesenchyme [51, 52, 53]. Another key difference lies in digit formation. In amniotes, digits form in a posterior-to-anterior sequence, through a paddle stage that is later sculpted by apoptosis in the interdigital regions [54]. In contrast, axolotl digits emerge sequentially in an anterior-to-posterior order, with each digit forming through localized outgrowth [46]. Despite these differences the fully patterned limb is remarkably similar. This raises the question whether the same underlying principles drive patterning in both systems, even if the specific outcomes may vary.

To inform the computational model, we imaged key genes involved in axolotl limb patterning at different developmental stages (Stages 44 to 47) using light-sheet fluorescence microscopy (Fig. 3A,B,E). Whole-mount samples were reoriented in 3D, and a representative 2D section capturing spatial gene expression patterns was analyzed (Fig. 3C). Figure 3A shows the expression of the *Hox* genes *Hoxa9, Hoxa11*, and *Hoxa13*, which encode PI along the proximodistal (PD) axis of the developing limb [47]. These markers broadly correspond to future stylopod (*Hoxa9*), zeugopod (*Hoxa11* and *Hoxa9*), and autopod (*Hoxa13, Hoxa11* and *Hoxa9*) regions. We observed that *Hoxa9* was expressed broadly throughout the mesenchyme from stage 44 onward. *Hoxa11* was expressed from the middle portion of the limb bud to the distal tip from stage 45 to stage 47. *Hoxa13* first appeared at the distal tip around stage 45.5 and remained restricted to the autopod region through subsequent stages. These results match what was observed by others [55]. We imaged mRNA expression associated with distal cues including *Wnt3a, Wnt5a*, and *Fgf8* (Fig. 3B) [53, 52, 51]. *Wnt3a* expression is localized to the limb epithelium. *Wnt5a* is expressed in both the epithelium and the mesenchyme at the distal tip. *Fgf8* is first detected at stage 45 in the distal mesenchyme and overlaps spatially with *Wnt5a*, though it emerges slightly earlier. All channels are shown together in composite images in Fig. 3D. The presence of similar expression domains in axolotls supports the idea that key signaling pathways are conserved, even in the absence of a morphologically distinct AER [56]. In Figure 3E, we imaged the expression of *Sox9*, a transcription factor marking pre-cartilaginous condensations. By Stage 48, distinct skeletal elements become visible, including the humerus (h), radius (r), ulna (u), and digits (d1, d2), providing a spatial reference to validate the GPP model predictions. Earlier stages (44-47) show the gradual emergence of these elements, with the humerus visible by Stage 45-45.5 and the radius and ulna beginning to appear by Stages 46-47.

**Figure 3.**
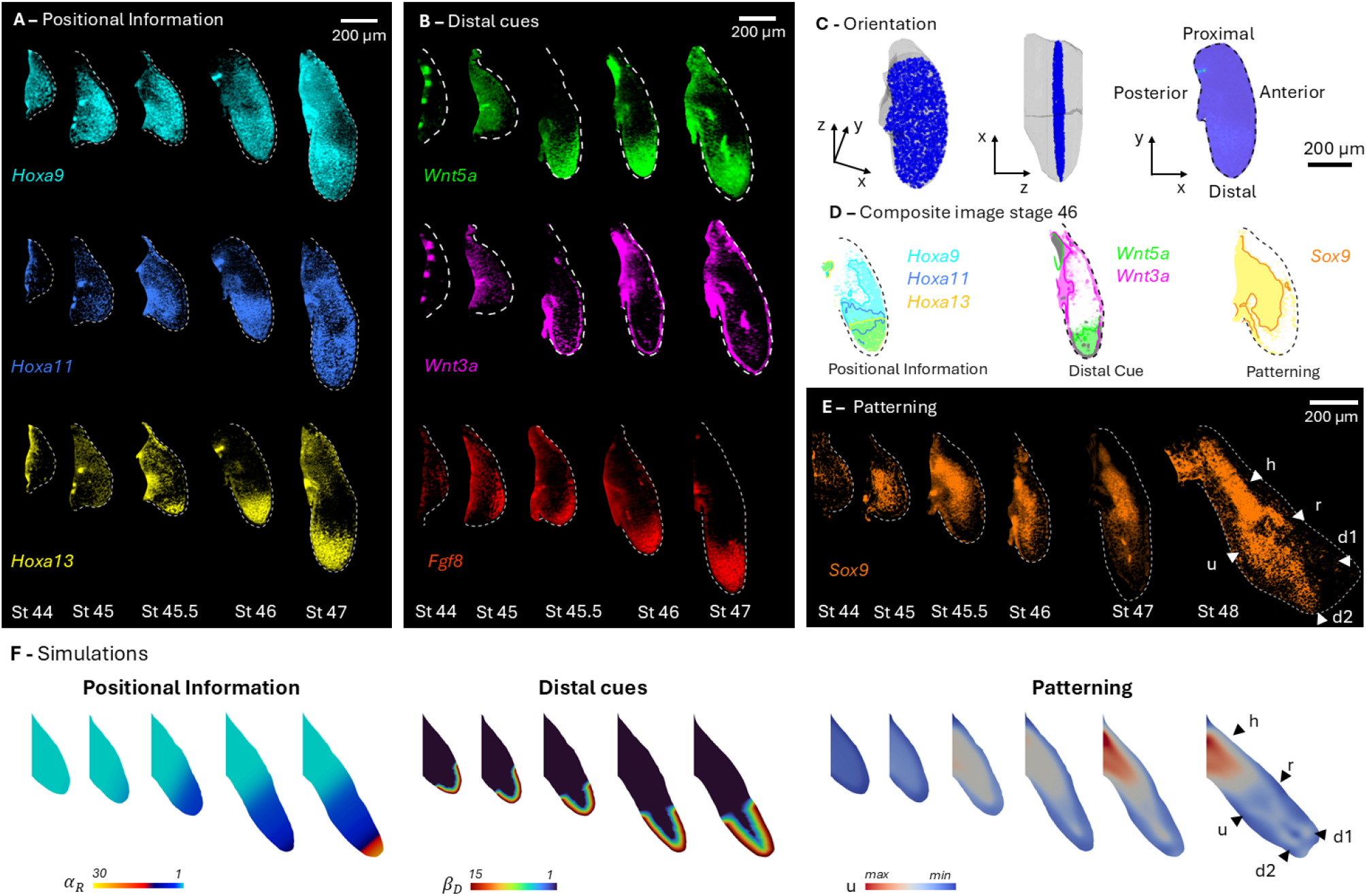
Experimental imaging and computational prediction with the GPP model of axolotl limb patterning. (**A**) Fluorescence imaging of *Hoxa9* (top row), *Hoxa11* (middle row), and *Hoxa13* (bottom row) at successive stages of axolotl limb development. These genes serve as markers of proximal-to-distal PI, and inform the spatiotemporal map of *α*_*R*_ in our simulations (**F**, left, “Positional Information”). (**B**) Fluorescence imaging of *Wnt5a* (top row), *Wnt3a* (middle row), and *Fgf8* (bottom row) key markers of the distal cue, and inform the spatiotemporal map of *β*_*D*_ in our simulations (**F**, center, “Distal cues”). (**C**) Orientation reference for the 2D plane (in blue) from the 3D limb imaging (in gray) shown in panels A, B, and E. (**D**) Composite representation of positional information (from *Hoxa* genes), distal signaling cues (from Wnt genes), and skeletal patterning at developmental stage 46. (**E**) Fluorescence imaging of Sox9, a marker of cartilage condensations, across developmental stages. At stage 48, skeletal elements are labeled: h (humerus), r (radius), u (ulna), d1 (first digit), d2 (second digit). (**F**) GPP model simulation results on the experimental limb geometry. The inputs *α*_*R*_ and *β*_*D*_ are informed by gene expression data (**A** and **B**); the output variable *u* predicts the spatial pattern of skeletal condensations (**E**).

Using the experimentally informed geometries and spatial inputs, we ran numerical simulations of the GPP framework to predict sites of skeletal patterning across developmental stages (Fig. 3F). In these simulations, *α*_*R*_ and *β*_*D*_ were assigned to reflect the distributions of *Hox* gene expression and distal signaling factors, respectively. Similar to the images in Fig. 3A, the segments’ identities are acquired progressively: the stylopod region (cyan) is established from the start, followed by the emergence of the zeugopod domain (blue) around Stage 45.5, and finally the autopod (orange) around the end of Stage 46 to Stage 47. Distal cues are applied at the distal tip from as early as Stage 44-45 and persist throughout the limb development. The resulting output variable, *u*, represents the skeletal element predicted. The model recapitulates the progressive emergence of distinct limb segments, beginning with a central condensation corresponding to the stylopod around Stage 45.5, which later bifurcates into two elements by Stage 46-47, followed by a second bifurcation that gives rise to additional elements aligned with digit positions observed in Sox9 expression at Stage 48. The spatial correspondence between the simulation output (Fig. 3F, right) and experimental marker expression (Fig. 3E) demonstrates that the underlying developmental logic captured by the GPP model is conserved and extensible beyond mouse limb development, without requiring structural modifications to the model itself. The role of AER signaling and PI is further explored in supplementary simulations (Fig. S1).

## 4 Discussion

### 4.1 Growth, Turing patterns and positional information in whole limb patterning

In this work, we introduce the Growth-Processing-Propagation (GPP) framework to provide a numerical tool for predicting patterning during limb formation. It is structured around three fundamental factors: 1) growth, which captures changes in size and shape; 2) processing, which includes gene regulatory interactions and their effects, such as reaction terms; and 3) propagation, encompassing morphogen signaling, cell-cell signaling, diffusion, etc. (Fig. 1A).

In vivo, these mechanisms operate through gene regulatory networks which represent the network of genes that determine cell fate [57]. For digit patterning specifically, researchers have identified several promising molecular candidates that can be modeled through RD systems, including the Sox9-Bmp-Wnt feedback loop [17, 58] and the TGF-*β*-Bmp loop [59]. Similar to previous models that assume that PI directly modifies molecular kinetics, our approach alters the weight of processing and propagation relative to growth using PI [18, 17]. The spatial and temporal continuity allows a systematic investigation of how different segments of the limb interact, respond to perturbations, and influence one another over time (Fig. 2C,D). We assess the removal and modification of the AER and PD cues and demonstrate altered patterning (Fig. 2A). This perspective also allows to explore the timing of PI emergence (Fig. 1B).

In the GPP framework, we use growth as the reference timescale, rather than reaction or diffusion, as in previous models [26, 60]. We chose this approach because growth is an observable process that can be quantified, whereas the timescales of reaction (processing) and diffusion (propagation) are more difficult to directly measure. These timescales can be understood as indicating which aspect of the system, growth, processing, or propagation, plays the dominant role at a given point in space and time. In our simulations the relative timescales, represented by *α*_*R*_ and *β*_*D*_, vary spatially over time.

Proximally, *α*_*R*_ = 1 and *β*_*D*_ = 1, indicating that processing, propagation, and growth all contribute to the emerging pattern. Distally, however, higher values of *α*_*R*_ and *β*_*D*_ (8 and 25) indicate that processing and propagation dominate over growth. This supports the assumption that, during autopod patterning, growth occurs on a slower timescale than signal processing and propagation, and can therefore be neglected in the reaction-diffusion equations. In contrast, the lower values observed in the zeugopod and stylopod suggest that growth is more tightly integrated into the patterning process in these regions. This spatial variation highlights the need to explicitly model growth through the convective term when addressing the entire limb. As illustrated in Fig. 5A, B, and Fig. S2, S3, S4, the growth rate distribution can significantly influence the pattern when *α*_*R*_ and *β*_*D*_ are low, while higher values yield outcomes similar to those on static domains or on growing domains without the convective term.

While this framework highlights the importance of growth, its current formulation has limitations. The model assumes no intrinsic directionality in growth and no built-in regional biases, meaning all parts of the domain are treated as equally permissive to growth. Despite this simplified starting assumption, the GPP model generates anisotropic and inhomogeneous growth patterns as a result of the boundary conditions (Fig. 4B,C). However, studies have shown that growth in the developing limb has inherent directional biases and regional variations [31, 32, 33, 61]. Our preliminary tests with different growth rate distribution shows that the specification pattern is significantly impacted (Fig. S3). These results emphasize the necessity of including the convective term in the equations (Fig. S2, S3, S4), particularly in early stages where the growth dynamics are equally as important as the RD system, and the importance of incorporating biologically realistic growth dynamics in future work. Moreover, since our model is two-dimensional, it does not account for patterning along the dorsal-ventral axis, meaning that spatial interactions in the third dimension are entirely missing from the model. Future work could integrate a more detailed understanding of growth to better align with biological reality. Growth, patterning, and cellular differentiation are inextricably linked during development [62], and the GPP framework does not yet fully capture the complexity of these interactions.

**Figure 4.**
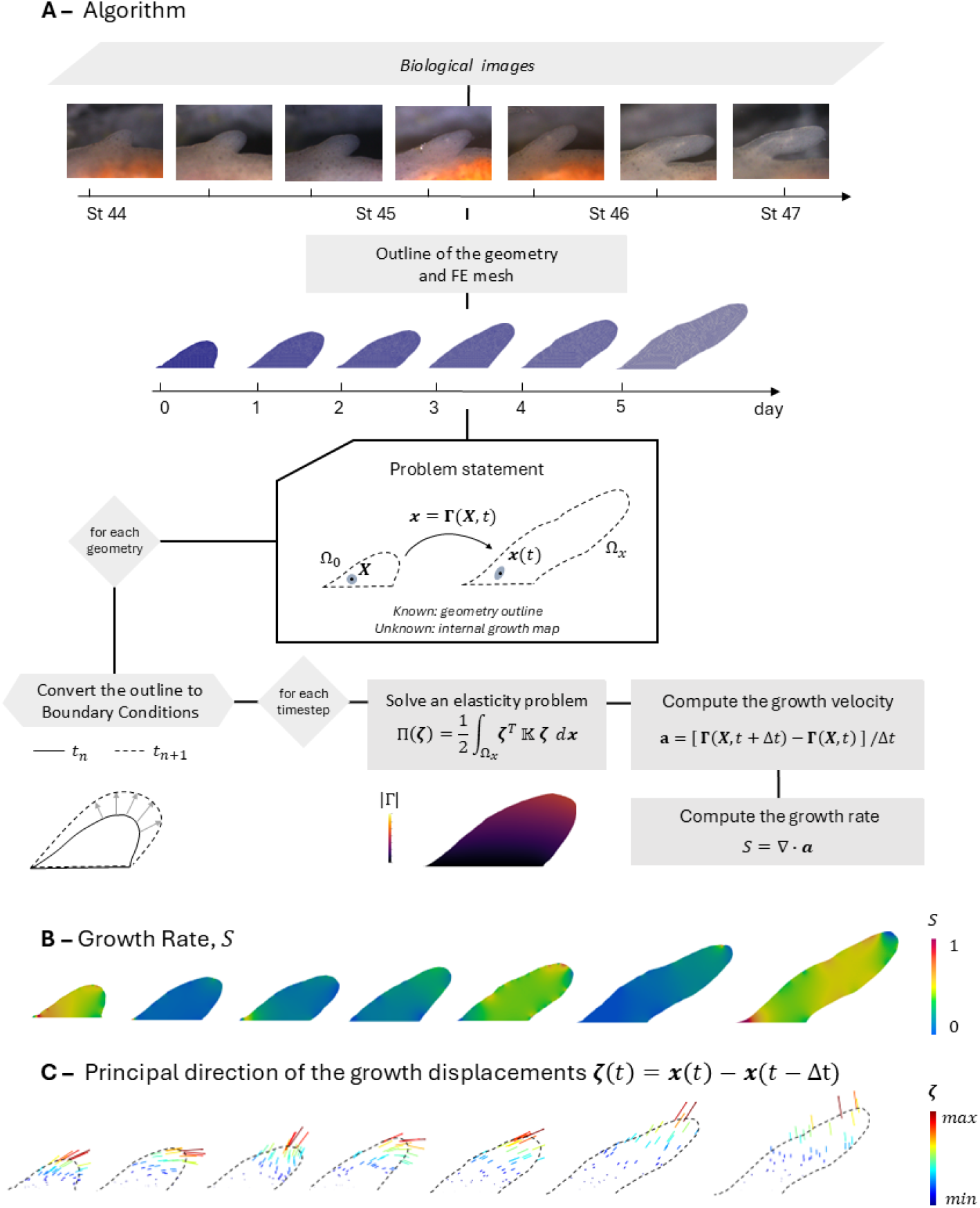
Numerical computation of growth rate from experimental images of developing axolotl limbs. (**A**) Stereo microscopy images were taken daily for a week from the same animal. The limb outline was manually extracted from these images and meshed using triangular linear elements. The material points **x**(**t**) represent the current configuration Ω_*x*_, which is related to their initial position **X** in Ω_0_ through the mapping function **Γ**(**X**, *t*). Each extracted outline was converted into boundary conditions, which were enforced in an elasticity problem to solve for the displacement field *ζ*. This displacement was then used to compute the growth velocity **a** and the growth rate *S*. (**B**) Computed growth rate *S* visualized across developmental time points, highlighting the non-uniform and dynamic nature of growth within the limb. (**C**) Principal directions of displacement vectors *ζ* at each node, revealing the anisotropic nature of tissue growth.

**Figure 5.**
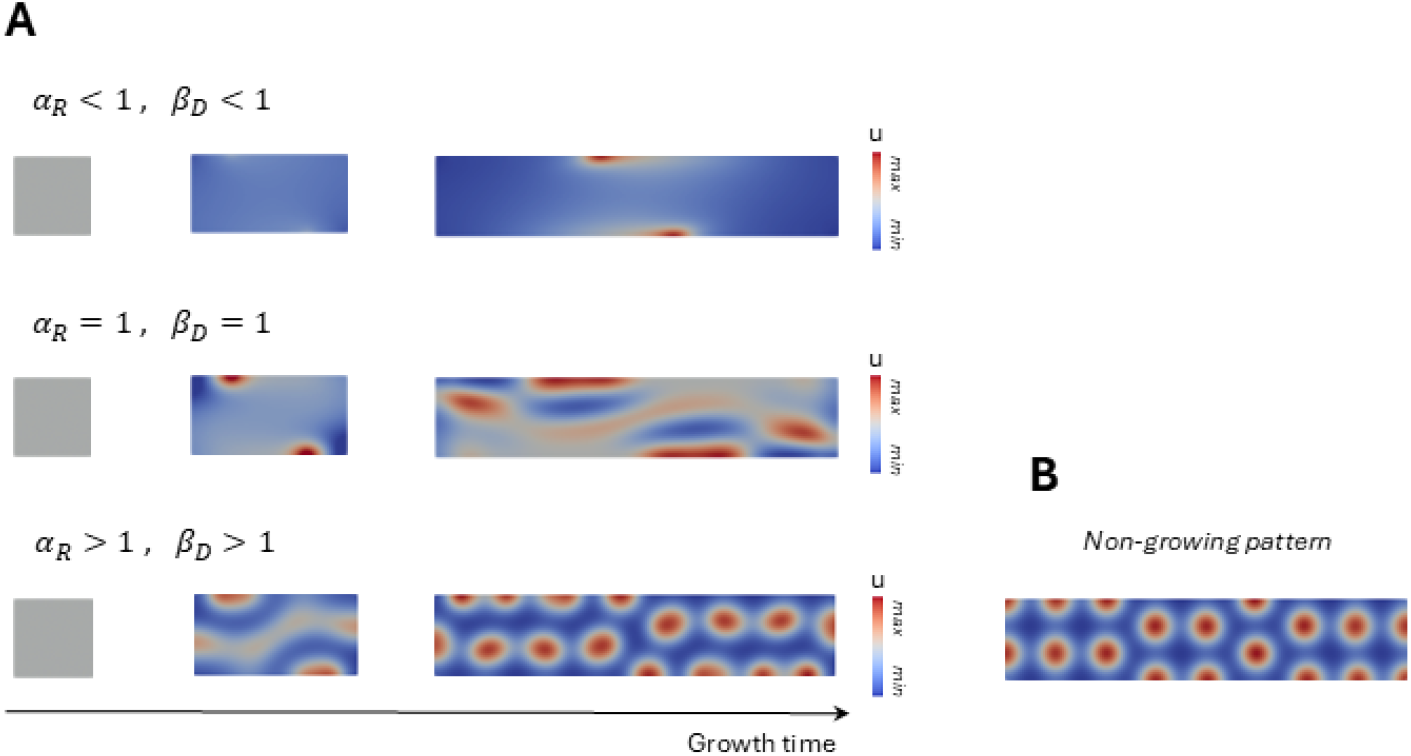
Growth-Reaction-Diffusion System. (**A**) Simulated patterns on a growing rectangle for varying *α*_*R*_ and *β*_*D*_. Results for an initial, intermediate and final domain size are shown. (**B**) Comparison with a non-growing rectangle (*α*_*R*_ *>* 1, *β*_*D*_ *>* 1)

### 4.2 The GPP framework from an evolutionary perspective

Limb development follows conserved patterning principles across tetrapods, yet morphological outcomes vary significantly between species. The concept of deep homology, which describes shared developmental mechanisms underlying diverse anatomical structures, suggests that common self-organizing principles shape limb formation across evolution [63]. The GPP approach provides a framework to model these shared processes by focusing on how spatial and temporal dynamics generate variability within a unified developmental model.

The GPP framework enables us to investigate homologous patterning mechanisms across species with different limb morphologies, such as amniotes and anamniotes (Fig. 1B, 3F). Previous studies have explored developmental variations by modifying spatial and temporal cues, focusing on skeletal elements (e.g. digits) or collection of discrete zones, such as active and frozen regions, with minimal coupling between them [64]. In contrast, GPP treats the whole limb as a continuous field. We can reproduce known changes in digit number (Fig. 2B) and, like existing studies, potentially link them to molecular regulators such as *Hox* genes [18]. However, the sensitivity of patterning to positional cues highlights the need for a more accurate spatial representation of these variations in the model (Fig. 2C and Fig. S5). Specifically, this would involve improving how imaging data is translated into model inputs to better capture the spatial variations in *α*_*R*_ and *β*_*D*_ distributions, allowing for more precise predictions of patterning outcomes [65].

Beyond capturing developmental variability, the GPP framework provides a tool for exploring evolutionary shifts in limb morphology. The evolutionary trajectories of mice and axolotls diverged over 365 million years, leading to distinct growth and patterning strategies. Yet, despite these differences, they can be studied within the same framework, enabling us to identify shared principles across tetrapods and explore a range of evolutionary changes. For example, features such as autopod elongation in bats [66], digit loss in tetrapods [14] or even limb regeneration properties [67] could be reexamined as a modulation of growth rate relative to processing and propagation. One of the most striking evolutionary transitions, the fin-to-limb shift, offers a compelling test case for this model [68]. In addition to studying changes like those in the AER, GPP provides a complementary perspective: could modifications in growth rate have also contributed to key structural changes? GPP serves not only as a predictive tool but also as a means to guide future experimental work, potentially revealing the molecular pathways underlying these transitions.

In summary, this study introduces the Growth-Processing-Propagation (GPP) framework as a conceptual and numerical tool to investigate how growth dynamics influence patterning in limb development. This perspective allows reinterpreting the interplay between RD systems and PI, offering a shared self-organizing mechanism for all limb segments. Beyond its developmental applications, the framework opens new avenues for exploring evolutionary transitions. Future work will extend the model to include realistic anisotropic inhomogeneous growth to further bridge the gap between theoretical models and biological complexity.

## 5 Materials and Methods

### 5.1 Growth

#### 5.1.1 Theory on growth

Biologically, growth results from a combination of cell proliferation, changes in cell size, cell migration, and production of extracellular matrix. These complex processes lead to both expansion of tissue volume and changes in tissue shape. These are approximated through an elastic deformation driven by the displacement of the limb profile. This deformation allows us to estimate a continuous growth rate from time-series images of developing limb. As the limb outline changes shape during development, the movement of the tissue, characterized by the trajectories of material points is calculated.

The initial position of a tissue element is defined in a two-dimensional domain using the Lagrangian coordinates **X** = (*X*_1_, *X*_2_). The position of a tissue material point is expressed by the bijective function **Γ**

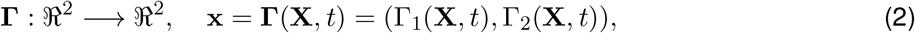

where **x** current spatial position at time *t*, with **Γ**(**X**, 0) = **X** and **Γ**^*−*1^(**x**, *t*) = **X**. We consider the deformation of the tissue entirely due to growth, and we deduce **Γ** using experimental data (see next section). The growth velocity at the current configuration is given by

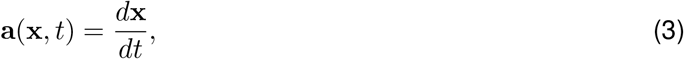

where the function **a**(**x**, *t*) is considered to be known and twice continuously differentiable. We define the scalar

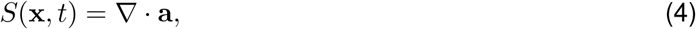

which measures the growth rate, i.e., the rate of area change per unit area.

#### 5.1.2. Derivation of growth from experimental images

To compute the growth rate from experimental images, the growing tissue is treated as an elastic continuum subjected to boundary displacement conditions. By constraining the boundary to match experimental outlines at different time points, we estimate how each point in the tissue moves over time and consider the displacements of the tissue solely due to growth. Experimental data providing the outlines of the geometry at multiple time points were used to deduce both (Fig. 4A). For the mice case, the outline was provided in [36], while for the axolotl, the outline was manually extracted from images using Fiji [69].

To compute **Γ**, the quadratic elastic energy function Π(***ζ***) is minimized with ***ζ*** being the displacement equal to ***ζ***(*t*) = **x**(*t*) − **x**(*t* − Δ*t*),

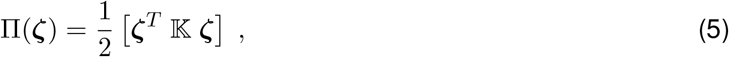

where 𝕂 is the growth rigidity matrix, which characterizes how tissue responds to boundary expansion. In the finite element implementation, it corresponds to the stiffness matrix. We assume a constant, homogeneous, and isotropic growth rigidity. This means we treat all regions of the tissue as equally responsive to growth. We recognize that this a simplification, as developing limbs exhibit spatial variations in proliferation rates and growth directions [31]. The minimization process finds how tissue expansion distributes most evenly throughout the growing domain while maintaining tissue continuity. Despite assuming homogeneous growth rigidity, this approach generates growth patterns that are non-homogeneous, anisotropic, and dynamic over time (Fig. 4B, C).

Numerically, the current position **Γ** is computed by using a finite element discretization [70] and constraining the displacements of the boundary nodes to match the experimental outlines.

There are as many geometries as the number of time steps. To obtain the outline for each time step, the geometry is linearly interpolated between experimental data points (Fig 4A). Then, Π(***ζ***) is minimized for each time step to deduce growth displacement, ***ζ***(*t*), which can be related to the growth velocity as

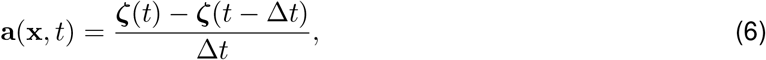

where ***ζ***(*t* − Δ*t*) is the displacement of the previous time step, and Δ*t* is the size of the time step.

### 5.2 Reaction-diffusion equation on growing domain

The reaction-diffusion equation in a growing domain, given in (1), involves a two-species vector **c** ∈ ℝ^2^ with a Schnakenberg reaction term **R** [71], and is solved using the finite element method. For more details on the governing equation, please refer to the Supporting text.

#### 5.2.1 Definition of the timescales

For simplicity, the equations are presented for a single species, but the non-dimensionalization process can represent a system that governs the behavior of multiple species. The process of non-dimensionalization captures the interactions and interdependencies within the system.

#### 5.2.2 Non-dimentionalization

##### Definition of the timescales

To non-dimensionalize the system, we define 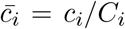 and 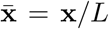, where *C*_*i*_ is a reference concentration for the *i*^th^ chemical species and *L* is a characteristic length, which we choose to be the initial length of the domain *L*_0_. The reaction rate *ω*, characteristic of the kinetic scheme, is used to non-dimensionalize the reaction term 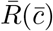. The diffusivity of each chemical species is denoted as *D*_*i*_, and we define *D*_1_ = max {*D*_*i*_}. The non-dimensionalized diffusivity is introduced as 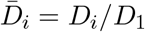.

Two timescales corresponding to the two elements on the right of (1), *T*_*D*_ and *T*_*R*_ for diffusivity and reaction kinetics, respectively are introduced:

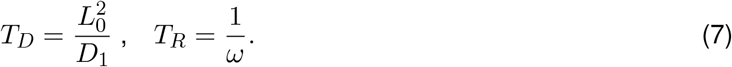

Here, the characteristic timescale *T*_*D*_ represents the time it takes for diffusion to spread across a distance equal to the characteristic length scale *L*_0_. The timescale *T*_*R*_ refers to the duration of a chemical reaction. It is typically measured as the time it takes for a change in the concentration of reactants to occur.

The characteristic timescale *T*_*G*_, which is associated with domain expansion due to growth, is defined as

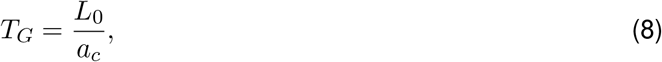

where the characteristic growth velocity

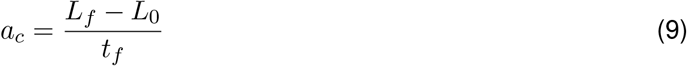

has been introduced. Here, *t*_*f*_ corresponds to the final time, at which the domain reaches the final length *L*_*f*_. Rewriting the growth timescale in terms of the final strain 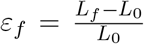 and using the relation *a*_*c*_ = *ε*_*f*_ *L*_0_*/t*_*f*_, results in *T*_*G*_ = *t*_*f*_ */ε*_*f*_, which provides an alternative interpretation for *T*_*G*_ as the time required to achieve a specific strain in the domain.

##### Time non-dimentionalization

Any of the three timescales, *T*_*G*_, *T*_*R*_ or *T*_*D*_, could be used to non-dimensionalize the time variable. Because the characteristic time of growth is the only experimentally measurable quantity in our problem, we use it to non-dimensionalize the time variable. Introducing *τ* = *t/T*_*G*_ into (1), we obtain

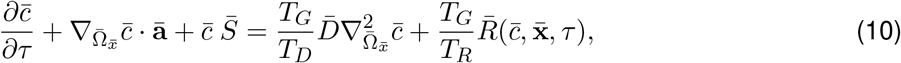

where 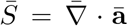 with ā = **a***/a*_*c*_ and *a*_*c*_ = *L*_0_*/T*_*G*_. The non-dimensional time *τ* can be related to the apparent strain as

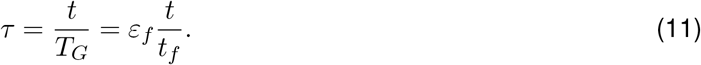

Dropping the bars and reverting *τ* back to a non-dimensional *t* to simplify notation, results in

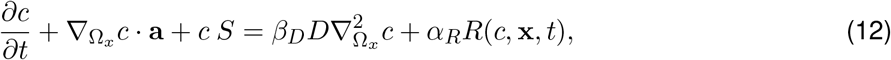

where the parameters *β*_*D*_ and *α*_*R*_,

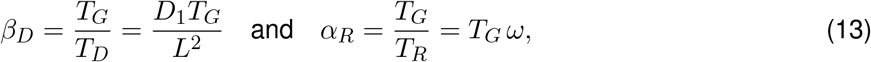

have been introduced.

One advantage of this approach is that the explicit knowledge of *T*_*G*_ enables straightforward conversion of nondimensional results back into dimensional units, allowing direct comparison with experimental data.

##### Interpretation of the parameters *α*_*R*_ **and** *β*_*D*_

The parameters *α*_*R*_ and *β*_*D*_ serve as indicators of the relative importance of reaction and diffusion compared to growth.

When *α*_*R*_ and *β*_*D*_ are less than one, growth dominates. In this regime, the pattern is strongly shaped by the growth rate distribution, through the convective term *S***c**, which represents the morphogen transport caused by tissue expansion due to growth (as shown in the Supplementary Information). When both parameters are close to one, all processes, reaction, diffusion, and growth, proceed on comparable timescales, yielding patterns shaped by the interplay of all mechanisms. When *α*_*R*_ and *β*_*D*_ exceed one, reaction and diffusion dominate over growth, The spatial pattern is governed primarily by the intrinsic properties of the reaction-diffusion system, and growth has negligible influence, producing outcomes similar to those observed in static (non-growing) domains (Fig. 5).

In practical terms, these parameters provide a framework for interpreting and estimating dimensional timescales. For example, in axolotl limb development, experimental data indicate a final apparent strain *ε*_*f*_ = 3.95 over a period of 12 days. This implies a characteristic growth timescale *T*_*G*_ = *t*_*f*_ */ε*_*f*_ ≈ 3.03 days. Based on the range of *α*_*R*_ values used in our simulations (from 1 to 30), the corresponding reaction timescales *T*_*R*_ = *T*_*G*_*/α*_*R*_ span from 3.03 days down to 0.101 days. Similarly, the diffusion timescales *T*_*D*_ = *T*_*G*_*/β*_*D*_, with *β*_*D*_ up to 15, span from 3.03 days down to 0.202 days. These estimates demonstrate how the non-dimensional framework allows re-dimensionalization of the system and facilitates interpretation across biological contexts.

### 5.3 Numerical Simulations

The governing equation (1) is discretized in space using the finite element method and in time with the implicit midpoint rule. The numerical implementation is performed in MATLAB [72] in an in-house code. Linear triangular elements with four-point Gauss quadrature are used to mesh the domain. The number of time increments is given by *ε*_*f*_ */*Δ*t*, with Δ*t* the time step. Homogeneous Neumann boundary conditions are applied. The function **R** follows Schnakenberg kinetics [71] with model parameters *a, b*, and *d* that ensure pattern formation (see supplementary text for details and refer to Table S1 for parameters used).

Simulations were initialized with random perturbations (10% variation) around the steady state defined by **R**(**c**) = **0**. For mouse and axolotl limb models, an epithelial layer was included at *t* = 0 with **c**(0) = **0** (Fig. 1, 2, 3). The effect of this initial condition is further explored in Fig. S6.

### 5.4 HCR-FISH and whole-mount hybridization imaging

Hybridization Chain Reaction Fluorescence In Situ Hybridization (HCR-FISH) was performed in whole-mount developing limb following the protocol provided by Molecular Instruments and as described by [73], without modification. Tissues were fixed in 4% paraformaldehyde, dehydrated in a methanol series, and stored at -20^*°*^C until use. Rehydrated tissues were treated with proteinase K, post-fixed, and hybridized with probes overnight at 37^*°*^C. After probe washes, HCR amplification was carried out using snap-cooled hairpins, followed by mounting in 1.5% low-melting agarose in glass capillaries and refractive index matched in *EasyIndex* (*LifeCanvas Technologies*).

Light-sheet imaging was performed on a Zeiss Z.1 microscope at 20X with dual side illumination. Refractive index matched, arose-embedded samples were imaged in EasyIndex. Image stacks were denoised in Zen Blue, and post-processing (rotation, cropping, and brightness/contrast adjustment) was conducted in Fiji and Matlab.

## Supplementary Information

### Supplementary figures & tables

**Figure S1.**
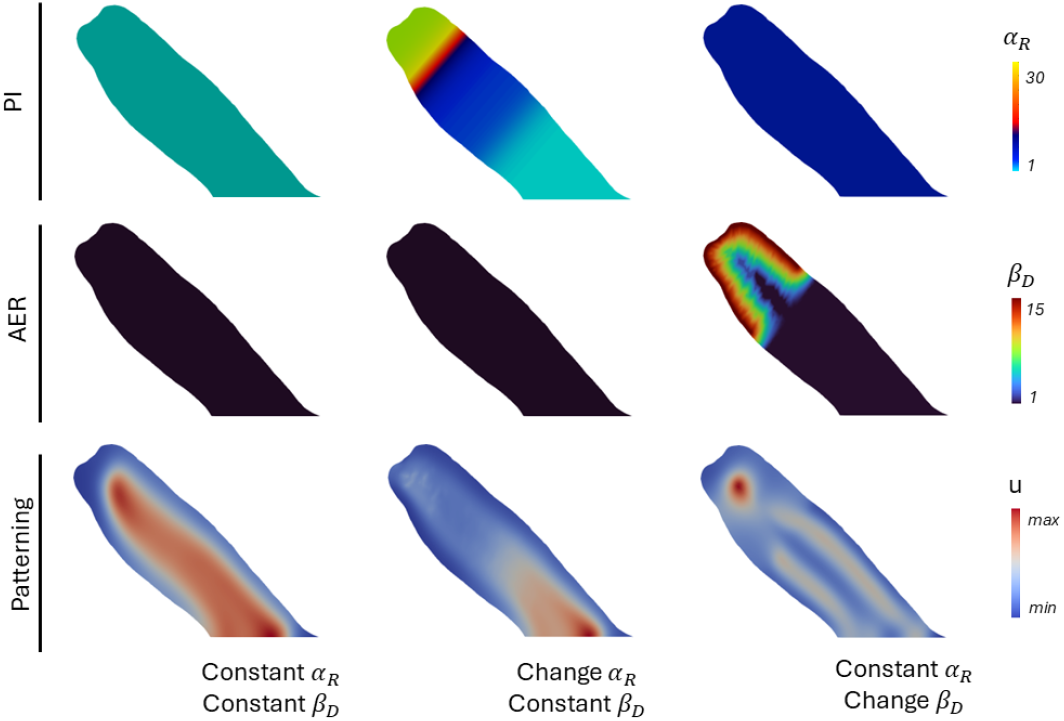
Effect of the distribution of *α*_*R*_ and *β*_*D*_ on the axolotl limb. The first column represents the framework without incorporating positional information (PI) or the apical ectodermal ridge (AER), implemented through homogeneous distributions of both *α*_*R*_ and *β*_*D*_, resulting in a simple stripe pattern. The second column includes PI but excludes AER, modeled by a homogeneous *β*_*D*_ and a spatially varying *α*_*R*_, maintaining the proximal pattern while losing autopod features. The third column incorporates AER but excludes PI, implemented by a homogeneous *α*_*R*_ and a spatially varying *β*_*D*_, where PI takes a uniform value corresponding to the zeugopod. As a result, the model produces an extended zeugopod-like element.

**Figure S2.**
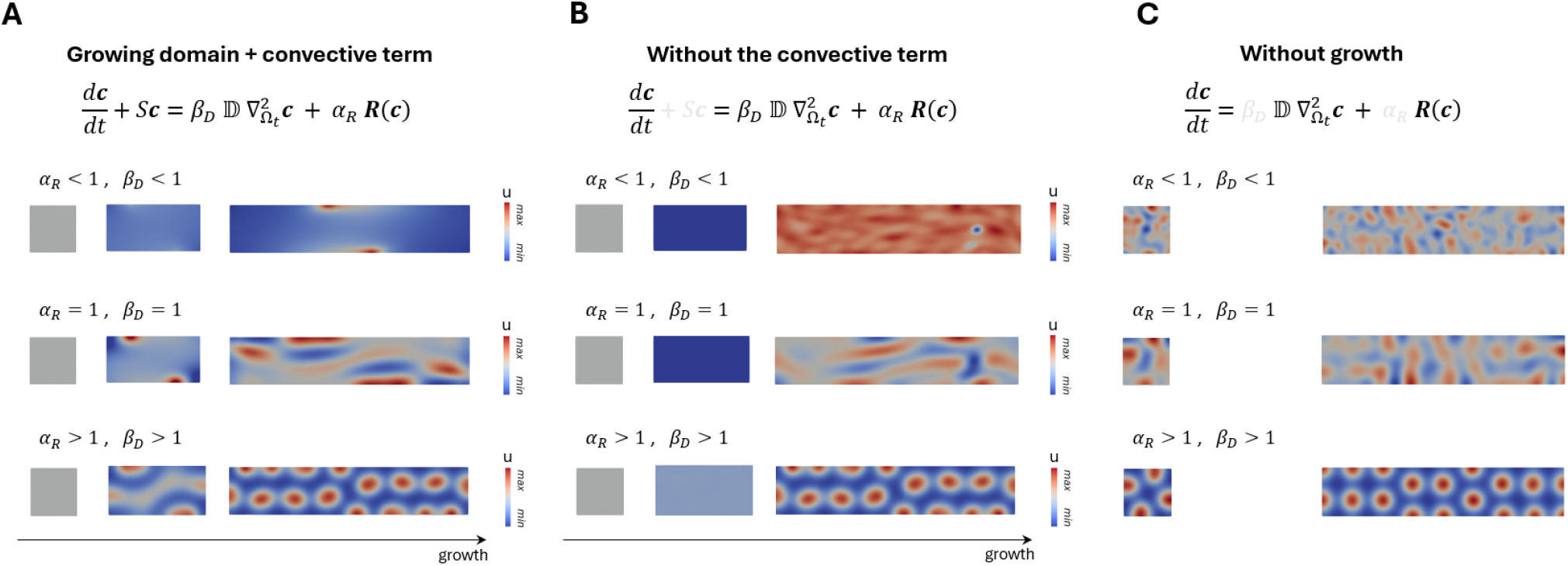
Effect of the convective term on pattern formation in growing domains. **(A)** Simulation on a growing rectangular domain including the convective term, as described in the main text. **(B)** Simulation on a growing domain without the convective term. Lower values of *α*_*R*_ and *β*_*D*_ show a noticeable impact of the convective term on the resulting pattern. **(C)** Simulation on a static domain (no growth), where the convective term is absent by definition. In this case, *α*_*R*_ and *β*_*D*_ lose their influence on the pattern’s shape, and all simulations yield the same final pattern. Higher values of *α*_*R*_ and *β*_*D*_ only accelerate the convergence of the pattern.

**Figure S3.**
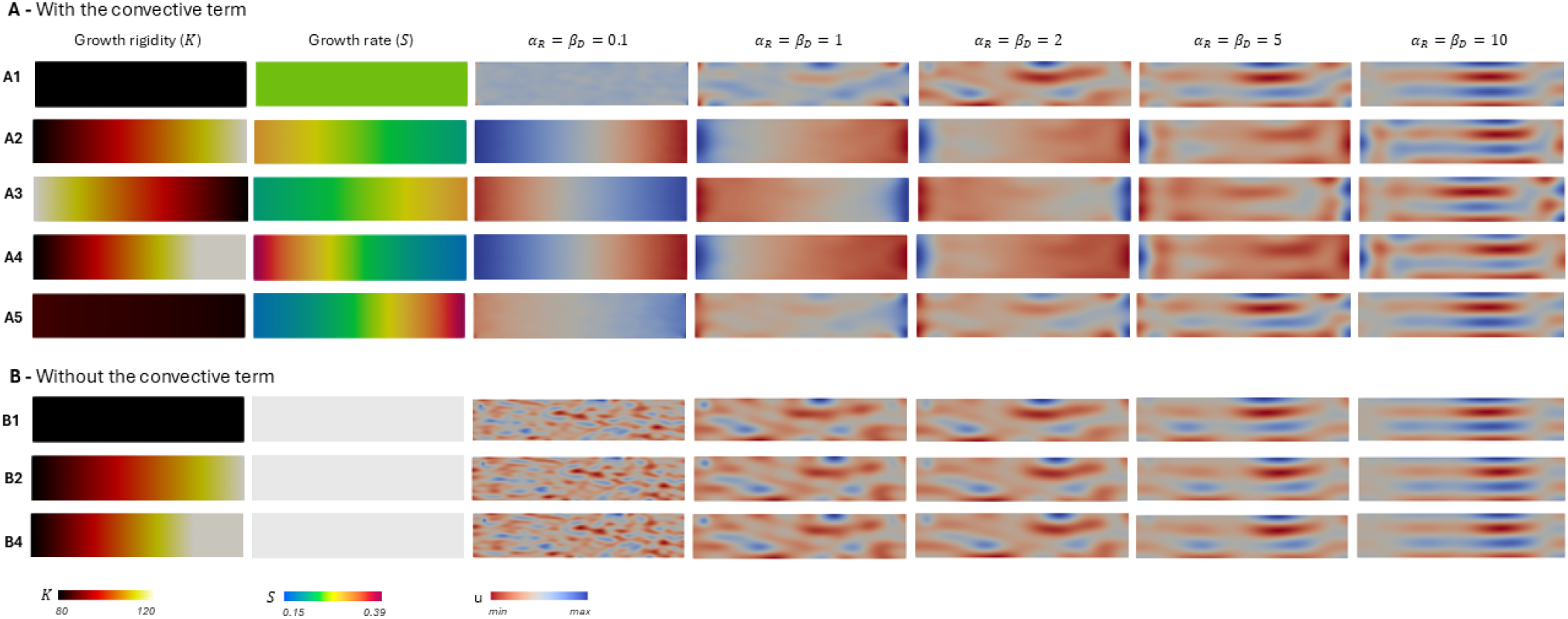
Effect of growth rate distribution *S* on pattern formation in growing domains. **(A)** Simulations on a growing rectangular domain including the convective term, as described in the main text. Each line corresponds to a different distribution of growth rigidity: A1 represents homogeneous rigidity, resulting in a uniform growth rate; A2 and A3 show a linear variation in rigidity, producing a linear gradient in growth rate; A4 and A5 implement a sigmoidal variation, generating a localized growth zone. Each simulation was run across a range of *α*_*R*_ and *β*_*D*_ values. At low values of *α*_*R*_ and *β*_*D*_, the spatial distribution of growth significantly affects pattern formation. In contrast, at high values of these parameters, reaction-diffusion dynamics dominate, leading to patterns that resemble those obtained in the absence of growth. **(B)** To assess the effect of mesh deformation alone, simulations were repeated without the convective term, using homogeneous (B1), linear (B2), and sigmoidal (B4) growth rigidity profiles. The results show that, even without the convective term, high *α*_*R*_ and *β*_*D*_ values lead to patterns comparable to those obtained with growth-free simulations, confirming the dominance of RD dynamics under these conditions.

**Figure S4.**
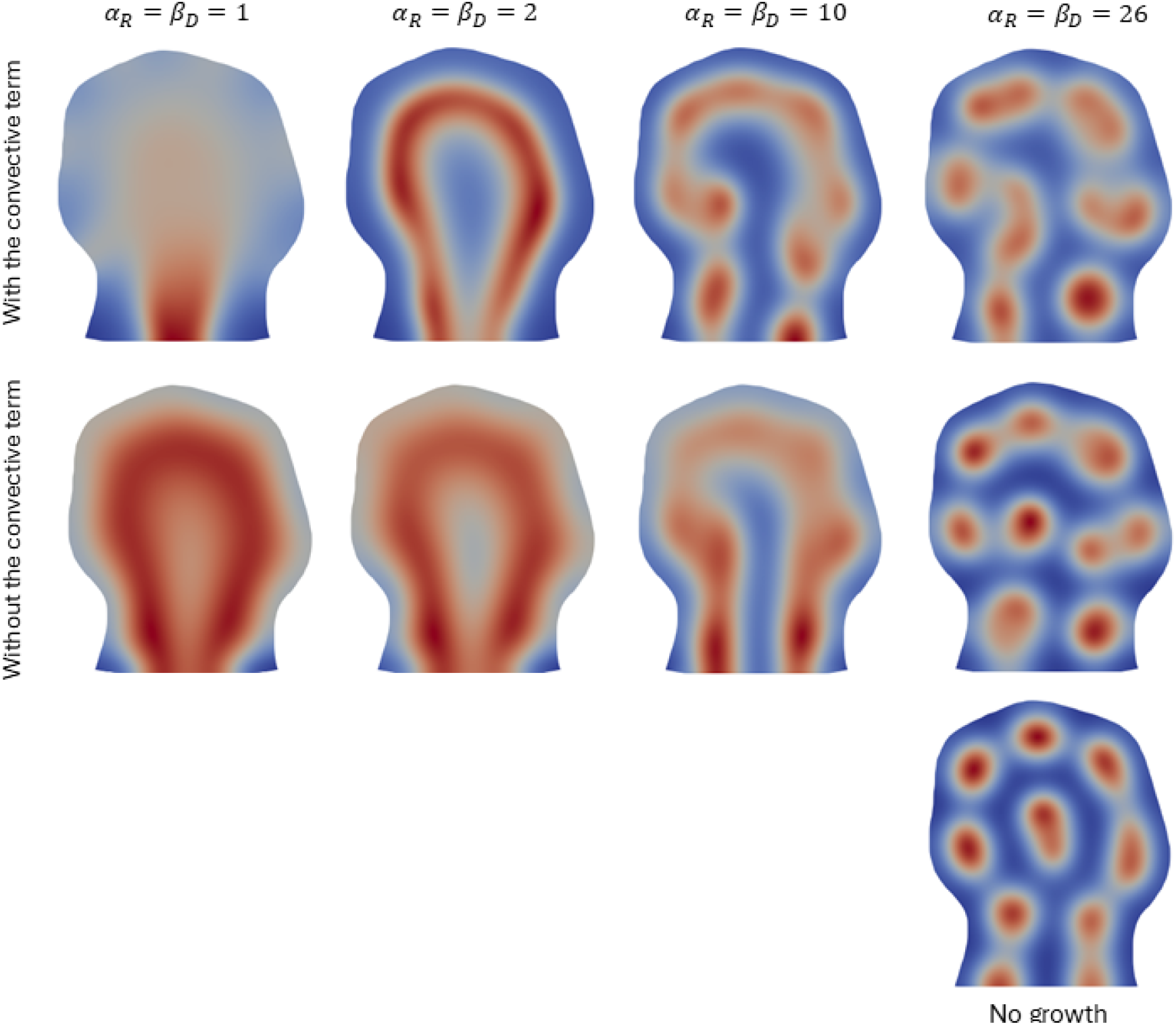
Effect of the parameters *α*_*R*_ and *β*_*D*_ on pattern formation. Simulations were conducted on a growing mouse limb bud domain, as described in the main text. Only the final geometry is shown. The first row presents results obtained with the convective term, and the second row without it, both across different combinations of *α*_*R*_ and *β*_*D*_, with parameters kept constant and homogeneous throughout the growth process. The third row shows patterns generated without growth, applied directly to the final limb bud geometry.

**Figure S5.**
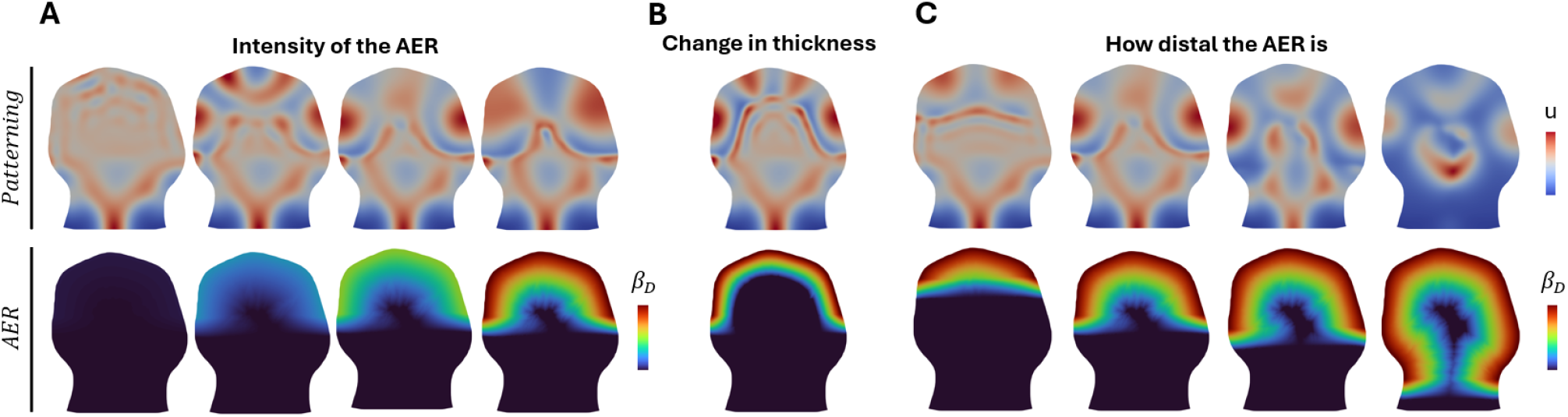
Influence of *β*_*D*_ distribution on limb bud patterning. (**A**) The intensity of *β*_*D*_, (**B**) Change in AER thickness, and (**C**) the proximodistal positioning of the AER influence patterning within the limb bud. The top row shows the resulting patterning in the limb, while the bottom row shows the corresponding AER. Color scales indicate the spatial variations in patterning (top row, *u*) and AER properties through *β*_*D*_ distribution (bottom row).

**Table: Parameter selection for patterning simulations**

For Fig. 5, the Turing space parameters are the same as those used for mice and axolotl. The values for *α*_*R*_ and *β*_*D*_ were set to 0.1 for low values and 10 for high values.

**Table S1.** Parameter values used for different cases. The parameters *a* and *b* correspond to the kinetic terms of the Schnakenberg system, as described in Eq. (S.11). The relative diffusion of *v* with respect to *u* (*d*) is seen in Eq. (S.10). The parameters *α*_*R*_, *α*_*R*1_, and *α*_*R*2_ represent the values of positional information shown in Fig. 1B and Fig. 3B. The parameters *l*_1_ and *l*_2_ define the transition positions between the stylopod and zeugopod, and between the zeugopod and autopod regions, respectively, relative to the final limb length. The parameters *a*_1_ and *a*_2_ control the smoothness of these transitions. The parameters *β*_*D*_ and *β*_*D*1_ indicate the AER values shown in Fig. 1B and Fig. 3B. Finally, *N* represents the number of elements used to define the thickness of the AER. AER parameters are explored in SUPP. S5.

| Parameters | Turing Space | | | $\alpha_R$ | | | | | | | $\beta_D$ | | |
| --- | --- | --- | --- | --- | --- | --- | --- | --- | --- | --- | --- | --- | --- |
| | $a$ | $b$ | $d$ | $\alpha_R$ | $\alpha_{R1}$ | $l_1$ | $a_1$ | $\alpha_{R2}$ | $l_2$ | $a_2$ | $\beta_D$ | $\beta_{D1}$ | $N$ |
| Mice | 0.47 | 2 | 113 | 1 | 2 | 0.13 | 0.03 | 8 | 0.45 | 0.03 | 1 | 26 | 30 |
| Axolotl | 0.47 | 2 | 113 | 1 | 5 | 0.25 | 0.05 | 35 | 0.75 | 0.05 | 1 | 15 | 12 |

**Figure S6.**
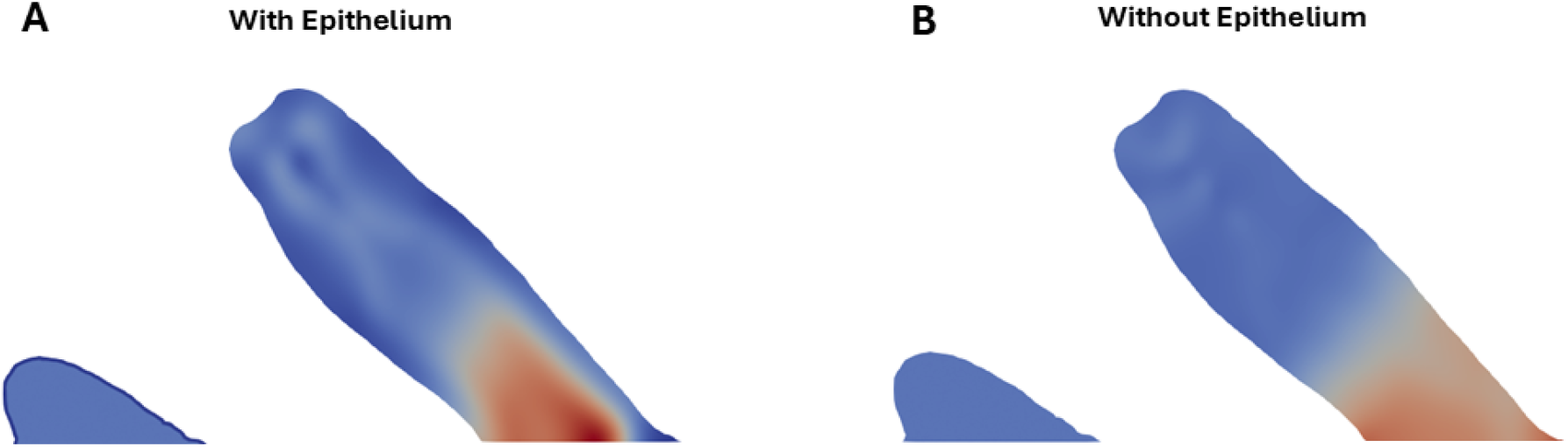
Effect of the epithelium on pattern formation. (**A**) Pattern formation with an epithelium boundary condition. The epithelium is modeled by setting *u*(0) = 0 and *v*(0) = 0 in the first two layers of elements. The rest of the elements are initialized at the steady-state solution (*u*_0_, *v*_0_), defined as **R**(*u*_0_, *v*_0_) = **0**. (**B**) Pattern formation without the epithelium. All elements are initialized at the steady-state solution (*u*_0_, *v*_0_). Both cases have different initial conditions at *t* = 0, without any additional specification on the epithelium for *t >* 0.

### Turing space through linear stability analysis

This section presents the derivation of the equations governing the Growth-Processing-Propagation system and describes the linear stability analysis used to identify the range of parameters that ensure pattern formation.

### Derivation of the reaction-diffusion equation in a growing domain

We consider a growing domain Ω_*x*_ enclosed by a surface *∂*Ω_*x*_. From the general conservation equation, the amount of material in Ω_*x*_ is equal to the rate of growth velocity of this material across *∂*Ω_*x*_ into Ω_*x*_, plus the material created in Ω_*x*_. For a species *ξ* with a concentration *c*, this can be expressed by

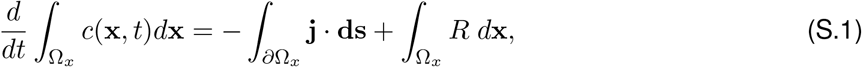

where **j** is the flux of material going through **ds** = **n** *ds* with **n** the normal of the surface element *ds*. The source of material is *R*, which can depend on the vector of species concentration *c*, the position vector **x**, and the time *t*. By applying the divergence theorem to the surface integral, the previous equation becomes

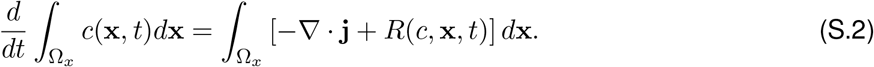

In order to evaluate the left-hand term, we use the Reynolds transport theorem

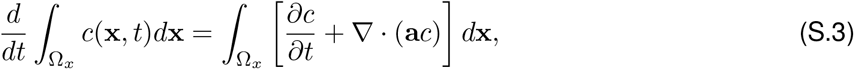

where **a**(**x**, *t*) is the growth velocity, and the evolution equation becomes

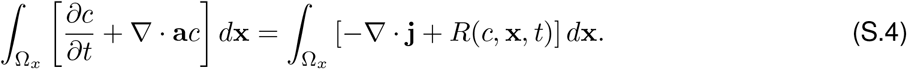

As this must hold true for any *t*, the integrands must be equal to each other. Therefore, we obtain the *local conservation equation* for *c* which is

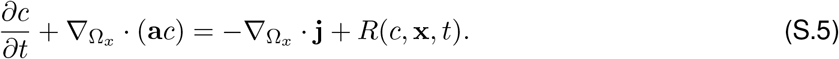

Here, **j** is a general flux transport vector, which can be diffusion or another process. We consider a classical diffusion problem,

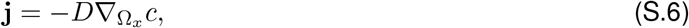

where *D* is the diffusivity at constant density. Equation (S.5) becomes

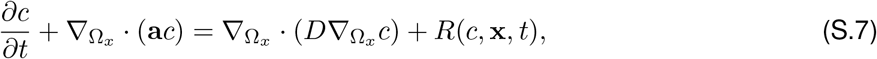

where *D* may be a function of **x** and *c*. The time-varying domain introduces two terms: ∇ *c* ·**a** and *c* ∇ ·**a**. The first represents the transport of material around the domain at a certain rate (dependent of the growth velocity **a**), in other terms it corresponds to the elemental volume moving with the growth velocity. The second one represents the dilution effect due to the local volume increase. The equation becomes

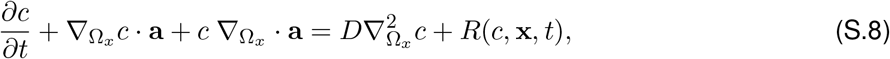

where *D* has been assumed to be independent of the spatial variable **x**. The final equation is

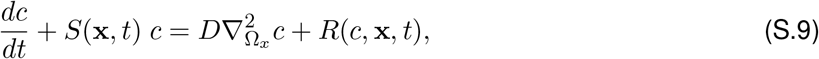

where *S*(**x**, *t*) is the growth rate.

### Linear stability analysis in a growing domain

We perform a linear stability analysis to determine the Turing space of parameters in a growing domain. First, we deduce the form of growth that yields a constant growth rate *S* in the linearized system. Then, we analyze its stability both with and without diffusion [60]. For pattern formation to occur, the system must be stable without diffusion and unstable when diffusion is present. These conditions yield four equations that define the set of parameters capable of generating patterns.

The linear stability analysis is then performed on a two-species system with **c** = (*u, v*),

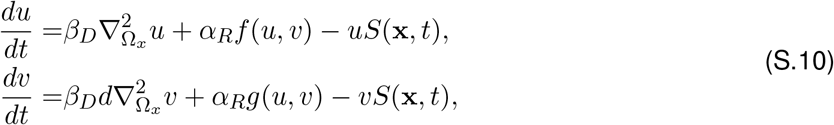

where *f* (*u, v*) and *g*(*u, v*) are the reaction terms for species *u* and *v*, respectively. The form of the functions follow the Schnakenberg kinetics [71]

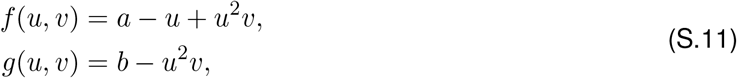

where *a* and *b* are model parameters. The ratio of their diffusivities is given by *d* = *D*_*v*_*/D*_*u*_. The parameters *α*_*R*_ and *β*_*D*_ are non-dimensional quantities. Our objective is to identify a homogeneous steady state that remains stable under small perturbations in the absence of diffusion, but becomes unstable when small spatial perturbations are introduced in the presence of diffusion.

The homogeneous steady state is defined as the solution of the system in (S.10) with both time and space independence. This solution can only be found if we assume that *S*(**x**, *t*) is a constant *S*. The homogeneous steady state can be determined by solving

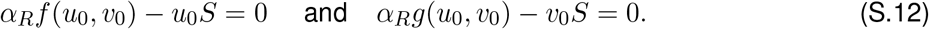

Introducing a small perturbation to the homogeneous steady state,

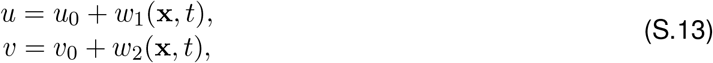

and substituting it into (S.10), the linearized system

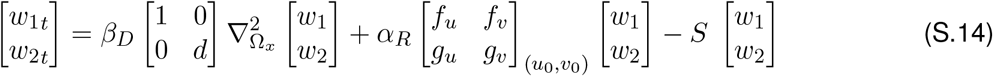

is obtained, where the subindex *t* indicates a time derivative, A is the Jacobian matrix of *f* and *g* evaluated at (*u*_0_, *v*_0_),

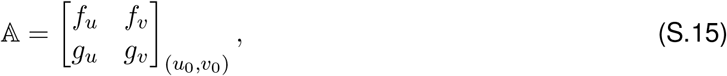

and

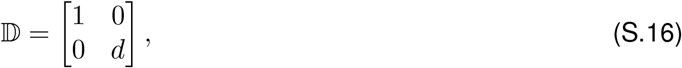

is the diffusivity matrix.

### Growth function for the linear analysis

To determine the linearized solution, the growth rate *S*(**x**, *t*) must be constant in both time and space. The goal is to identify a growth function **x**(*t*) = **Γ**(**X**, *t*) that yields a constant *S*. For simplicity, the analysis is restricted to one dimension.

An exponential growth function is first considered,

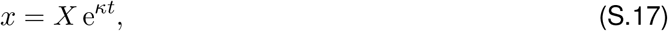

with growth velocity

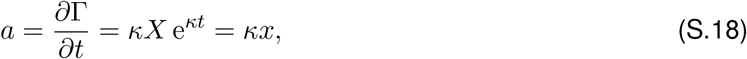

leading to a constant growth rate

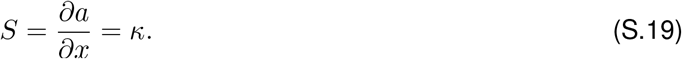

This exponential form maintains a constant *S*(*x, t*), satisfying the requirements for the linear stability analysis.

For comparison, a linear growth function is also examined,

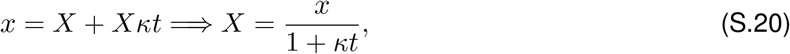

where *κ* is a known scalar. The corresponding growth velocity is

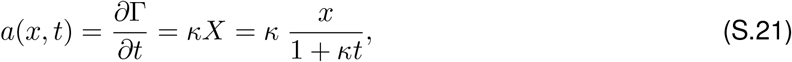

yielding a time-dependent growth rate,

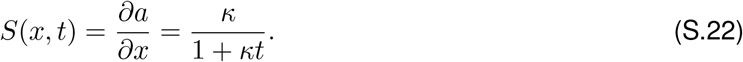

In this case, *S*(**x**, *t*) is unsuitable for the linearization. In the axolotl limb, experimental measurements suggest that growth is approximately linear in time, though spatially non-uniform.

### Linearized solution

We assume a general form for the solution to (S.10)

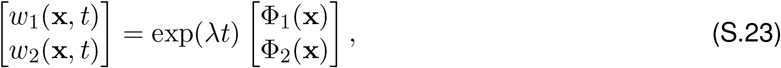

where *λ* is an Eigenvalue of *α*_*R*_𝔸, and Φ_1,2_(**x**) are the components of the Eigenvector of the spatial problem. Substituting this into the linearized equation (S.14) yields

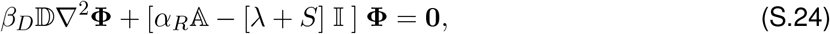

where 𝕝 is the 2×2 identity matrix and **Φ** is the vector [Φ_1_, Φ_2_]. The solution that satisfies (S.24) can be expressed as **Φ**_*m*_ = **y**_*m*_Φ_*m*_, where **y**_*m*_ is a constant vector and Φ_*m*_(**x**) is a scalar Eigenfunction of the Laplacian satisfying the boundary conditions. This implies that 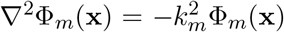 where *k*_*m*_ is the wavenumber corresponding to each Φ_*m*_. Therefore, (S.24) becomes

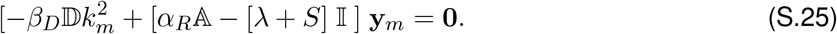

For non-trivial solutions, the determinant must satisfy

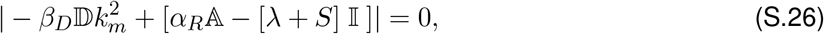

leading to the dispersion relation 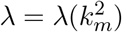. It can be written as

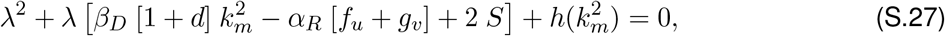

where

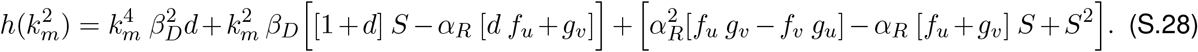

### Stability without diffusion

To determine the stability of the solution, we examine the sign of the real part of 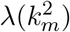. If *Re*(*λ*) *<* 0, the solution is stable (decreasing in time), and if *Re*(*λ*) *>* 0, the solution is unstable (increasing exponentially in time). Without diffusion (*k* = 0), the equation becomes

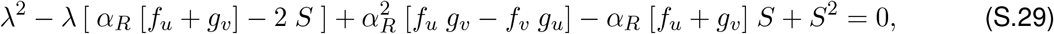

whose roots are

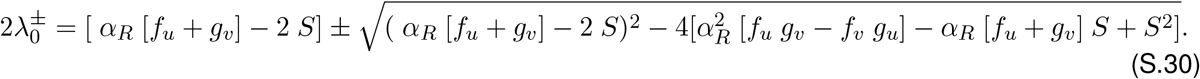

For 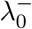, the condition for stability is

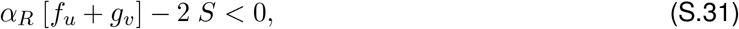

which can be rewritten in terms of the trace of A to obtain the first condition

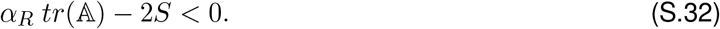

For 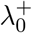 to be negative, we require

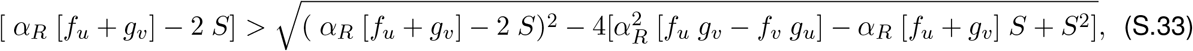

which can be written as resulting in

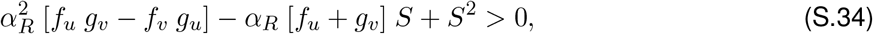

resulting in

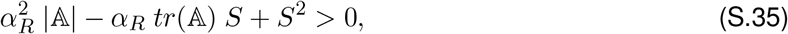

which is the second condition.

### Instability with diffusion

When considering diffusion in equation (S.27), the roots are

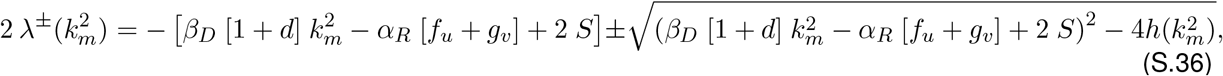

Given the condition for stability (S.32), necessarily

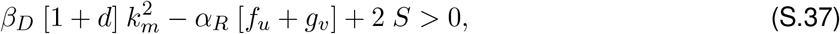

and therefore, 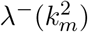 is always negative. To ensure instability, 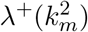 must be positive, which requires

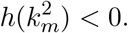

This leads to the condition

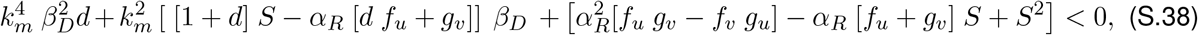

which, given the condition for stability (S.35), can be simplified to

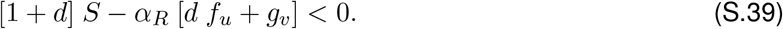

This relation provides a necessary condition but is not sufficient on its own. To establish a condition that is both necessary and sufficient, the minimum value 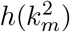 must be negative. This leads to the following condition

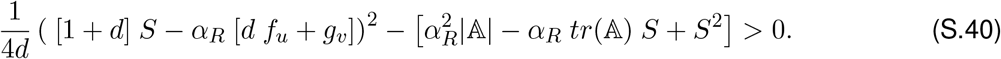

### Instability zone

The range of values 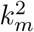 for which 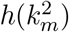 is negative is

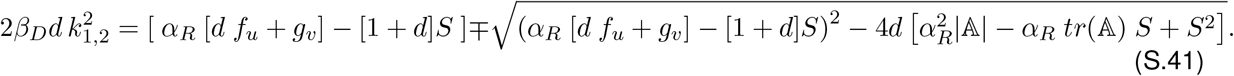

In two dimensions, in a rectangular domain *L*_*x*_(*t*) × *L*_*y*_(*t*) the condition becomes

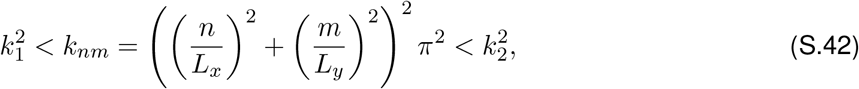

with *n* and *m* positive integers. It can be verified that for *S* = 0 (absence of growth), the standard conditions are recovered.

